# Biological foundation models illuminate annotation blind spots in evolutionarily divergent genomes

**DOI:** 10.64898/2026.05.15.724572

**Authors:** Toby B. Lanser, Sydney K. Caldwell, Gaspar A. Pacheco, Jessica W. Chen, Shahab Saghaei, Mariah Hassan, Meydan Kronrod, Duane R. Wesemann, H. Robert Frost

## Abstract

Chromosome-scale assemblies are increasingly available for non-model organisms, but functional annotation remains limited when deep evolutionary divergence erodes primary amino-acid sequence identity even though protein structural similarity can remain conserved. We present a hybrid annotation framework that decouples gene-model discovery from cross-species similarity assignment by combining Evo2-based *ab initio* prediction of exon–intron structures with ESM-2 protein-embedding-based structural similarity mapping. Applied to the sea lamprey, the framework derives high- or medium-confidence cross-species similarity assignments for 73,485 Evo2-derived translated protein models, including 35,395 high-confidence calls, and expands the deduplicated structural catalog to 31,286 loci, including 20,871 additions absent from the Ensembl baseline. A joint alignment–structure classification identifies 21,391 structurally supported catalog loci that a fixed human DIAMOND protein search does not confidently assign on its own, including 21,184 loci with no detectable human protein-sequence match and 207 loci with only low-confidence matches in the classical 20–30% amino-acid-identity twilight zone. These rescue-space totals describe catalog loci rather than validated one-to-one human-absent genes. In a single-cell RNA sequencing application, a stricter UTR-aware Ensembl+Evo2 reference improves gene recovery and expands the interpretable feature space of the lamprey immune compartment relative to the Ensembl baseline. This enables more resolved annotation of four transcriptionally defined immune cell states, including VLRA^+^-associated T-like and VLRB^+^-associated B-like programs together with oxidative iron-handling and iron-associated VLR-linked states. Together, these results show that structural protein signal often persists beyond the limits of pairwise sequence alignment and that an embedding-based annotation layer can extend that signal to improve downstream comparative and single-cell analyses in evolutionarily divergent genomes.

## 1 Introduction

Current genome annotation relies on “transitive annotation,” which transfers gene names and functions from model organisms to a new genome through pairwise protein-sequence alignment (Buchfink et al. 2015; Yates et al. 2020). At evolutionary distances beyond 500 million years (Myr), amino-acid identity in the primary sequence, the linear order of residues in a protein, often falls into the classical 20–30% sequence-similarity “twilight zone,” where pairwise alignments retain only weak residual signal and become difficult to distinguish from background similarity (Rost 1999; Iovino and Ye 2024). In that regime, alignment-based transfer workflows still depend on detectable amino-acid conservation, so similarity-based label transfer becomes increasingly incomplete as substitutions, insertions, deletions, and domain remodeling accumulate (Buchfink et al. 2015; Yates et al. 2020). As a result, failure to recover a convincing protein-sequence match in a deeply divergent genome does not imply absence of a bona fide protein-coding gene; instead, many loci remain labeled as “hypothetical” or “orphan,” limiting biological interpretation (Tautz and Domazet-Loso 2011; Vakirlis et al. 2020). Here, we distinguish taxonomically restricted genes from annotation orphans, which lack a detectable cross-species match under the search strategy at hand.

Recent sequence-only annotation models such as ANNEVO show that gene structures can increasingly be inferred directly from genomic sequence without external evidence (Zhang et al. 2026). In highly divergent genomes, however, accurate exon–intron structure still leaves the main comparative problem unsolved. Put directly, once pairwise alignment among translated proteins stops being informative, how do we derive plausible cross-species similarity assignments for newly predicted loci and determine whether those assignments improve downstream interpretation? The problem becomes sharpest in two settings: complete alignment loss and weak residual similarity in the classical twilight zone.

We address this bottleneck by decoupling gene-model discovery from cross-species similarity mapping using two classes of foundation models. First, **syntactic discovery** via Evo2 predicts exon–intron structures and coding coordinates from DNA syntax alone, including splice sites, codon usage, and promoter motifs, without requiring homology or transcriptomic evidence (Brixi et al. 2026; Stanke et al. 2006; Brŭna et al. 2021). Second, **semantic mapping** via ESM-2 protein embeddings places predicted proteins into a cross-species structural similarity space as a vector-search problem rather than string matching, leveraging the greater conservation of 3D protein structure relative to primary amino-acid sequence (Lin et al. 2023; Illergard et al. 2009). In practice, this step derives a provisional cross-species similarity assignment from the nearest structurally similar reference protein; it does not by itself establish one-to-one orthology for every call. We use embeddings when pairwise alignment of translated proteins no longer provides a clear assignment, because fold-level protein similarity can persist after amino-acid identity drops into the twilight zone. This framework therefore aims to recover structurally supported cross-species similarity assignments for predicted loci that lie beyond the practical reach of pairwise protein-sequence alignment alone and to make those loci usable for downstream comparative and single-cell analyses.

We apply this framework to the sea lamprey (*Petromyzon marinus*), an early-diverging jawless vertebrate separated from jawed vertebrates by about 550 Myr and with a highly repetitive genome (Smith et al. 2013). In the comparative panel used here, lamprey anchors the cyclostome branch; elephant shark, zebrafish, and human span cartilaginous, teleost, and mammalian gnathostomes; octopus serves as the bilaterian outgroup. The resulting annotation recovers immune- and developmental-linked similarity assignments, defines a vertebrate-biased structural cohort, and improves the single-cell reference for the lamprey immune compartment.

## 2 Methods

### 2.1 Data sources

Primary inputs were the larval sea lamprey (*Petromyzon marinus*) genome assembly and Ensembl annotation backbone, Evo2 *ab initio* gene predictions, and reference proteomes from human (*Homo sapiens*), zebrafish (*Danio rerio*), elephant shark (*Callorhinchus milii* ), and octopus (*Octopus sinensis*) for cross-species similarity mapping. The single-cell dataset comprised sea lamprey whole-blood 10x Genomics Chromium single-cell RNA sequencing (scRNA-seq) profiles from two PBS-injected control samples used for downstream reference evaluation. Whole-blood cells were collected from larval lamprey, enriched for live DAPI-negative cells by flow sorting, and sequenced on the Chromium GEM-X Single Cell 5′ v3 platform; Supplementary Methods S5 gives the experimental protocol, library construction, and Cell Ranger preprocessing. The lamprey baseline annotation was anchored to Ensembl release 110 on the larval assembly Pmarinus 7.0; Supplementary Methods lists the exact source files, versions, and preprocessing steps for each dataset.

### 2.2 Annotation and structural similarity pipeline

We ran the annotation pipeline in three phases. First, gene boundaries were predicted using evidence-based HMMs (BRAKER2) together with *ab initio* genomic language model inference from Evo2 on both strands (Brixi et al. 2026; Brŭna et al. 2021; Stanke et al. 2006). Second, predicted proteins underwent ORF-aware translation correction in a Biopython-based workflow, were embedded via ESM-2 (*N* = 19,850 human + *N* = 73,585 lamprey proteins) into a 1,280-dimensional structural space, and were matched to the human, zebrafish, elephant shark, and octopus reference proteomes using FAISS (Cock et al. 2009; Lin et al. 2023; Johnson et al. 2021). Third, translated protein models were binned into phylogenetic cohorts using cosine similarity between embedding vectors. The reference panel spans mammalian, teleost, and cartilaginous-fish gnathostome diversity while retaining octopus as a bilaterian outgroup. Throughout the manuscript, *S* denotes this embedding-based cosine similarity statistic, with larger values indicating greater structural similarity in ESM-2 space. Empirical thresholds were derived from empirical null distributions (vertebrate: *S* ≥ 0.9587; bilaterian: *S* ≥ 0.9745; see Supplementary Methods for full details).

For the Ensembl+Evo2 consensus annotation, loci were collapsed into named cross-species similarity groups using the final assignment fields carried through the pipeline. The `human_hit` field records the primary human-match term from the embedding-search workflow, and `final_name` stores the resolved display name when manual or rule-based reconciliation was needed.

To project cross-species similarity groups into phylogenetic structural space, each group was summarized by its cosine similarity to the octopus outgroup and to the three vertebrate anchors (elephant shark, zebrafish, and human). The octopus branch-shift statistic used in the transcription-factor analysis was defined as mean vertebrate similarity minus octopus similarity for a given similarity group or family.

### 2.3 Pathway and transcription-factor analyses

Pathway enrichment was performed with cameraPR (limma) using Reactome gene sets from msigdbr, mapped to lamprey via the human_hit human-match assignments (Supplementary Methods S6) (Ritchie et al. 2015; Dolgalev 2022). The ranking statistic was each lamprey gene’s maximum cosine similarity to the human reference embedding set, so enriched pathways represent groups of genes whose best structural matches to human are stronger than expected relative to the genome-wide background.

To assess regulatory remodeling without inflating broad Gene Ontology categories, we performed a separate genome-wide transcription factor analysis using the curated Lambert/Weirauch human TF census (Lambert et al. 2018). TF symbols were mapped onto the Ensembl+Evo2 consensus cross-species similarity assignments through `human_hit` and, when needed, `final_name`; family-level summaries were then computed from vertebrate mean similarity and the Octopus branch-shift statistic, defined here as mean vertebrate cosine similarity minus Octopus cosine similarity for the same TF group (Supplementary Methods S7).

### 2.4 scRNA-seq reference construction and rescue-space classification

For 10x 5′ data, reference design matters because gene-expression reads concentrate near transcript starts, first exons, and adjacent 5′ untranslated regions. Pure *ab initio* models often miss those captured regions when they predict only CDS coordinates, so the capture window can fall outside annotated transcript boundaries or compete with overlapping models. We therefore maintain two outputs throughout the manuscript. A broad **structural catalog** supports evolutionary and cross-species similarity analyses (31,286 deduplicated catalog loci, not 31,286 experimentally validated distinct genes), while a stricter **UTR-aware single-cell RNA-seq (scRNA-seq) reference** supports quantification. The scRNA reference starts from Ensembl’s UTR-annotated gene set (*N* = 13,114) and adds only non-overlapping Evo2 predictions that pass a stricter scRNA-specific filter (*S* ≥ 0.99; *N* = 17,244), yielding a 30,358-gene reference (Yates et al. 2020).

For the immune-subset marker analysis, marker genes were ranked by Wilcoxon differential expression across Leiden clusters and the top 25 markers per cluster were classified by annotation provenance (named Ensembl baseline, rescued cross-species assignment, anonymous Ensembl ID, or novel Evo2 model). We refer to the fraction of top markers assigned through rescued cross-species mapping as the **rescued-marker burden**. In parallel, we defined five hand-curated functional modules spanning T-like identity, B-like identity, iron handling, complement/innate signaling, and stress response, and summarized them at the cluster level with Scanpy gene-set scores (Wolf et al. 2018).

We also constructed a canonical alignment–structure classification on the 31,286-locus structural catalog. Loci with strong structural support (*S* ≥ 0.9587), where *S* is the embedding-based cosine similarity to the best matched reference protein, were partitioned into **alignment-supported** (≥ 30% amino-acid identity to the best matched human protein), **low-confidence human match** (*<* 30% amino-acid identity, corresponding to the classical 20–30% twilight zone), and **no detectable human protein-sequence match** (no measurable human protein hit under our sequence-search settings). This classification yielded both a measurable-hit subset for direct amino-acid-identity analysis and a broader rescue-space estimate that included all loci meeting the structural criterion regardless of whether protein-sequence alignment returned a weak hit or no hit at all. These rescue-space totals quantify how often the structural catalog escapes a fixed human protein search; they are not meant to imply the same number of biologically distinct human-absent lamprey genes (Supplementary Methods S3).

### 2.5 Animal care

*Petromyzon marinus* larvae (9.5–10 cm, 0.85–1.0 g) were obtained from the U.S. Geological Survey (USGS) Hammond Bay Biological Station, Michigan, and housed in temperature-controlled tanks with aerated filtered water at 17–21 °C, pH 7.5–8.3. All animal procedures were approved by the Brigham and Women’s Hospital Institutional Animal Care and Use Committee (IACUC) under protocol 2023N000131.

### 2.6 Cell preparation

Larval *P. marinus* were euthanized in tricaine methanesulfonate buffered in Tris-HCl (pH 8.0) until unresponsive to tail pinch. Blood was collected from the tail vein after decapitation, centrifuged at 300 × g for 5 min, and resuspended in 1 mL RBC lysis buffer (Invitrogen, cat. no. 00-4333-57). After 5 min at room temperature, cells were pelleted, washed with 1 mL of 0.67 × PBS + 1% FBS (L-FACS buffer), filtered through 40 μm mesh, and re-pelleted. Cells were stained for 20 min on ice with 2.5 μg/mL DAPI in L-FACS buffer, washed, and resuspended in 200 μL L-FACS buffer. DAPI-negative live cells were sorted on a FACSAria Phusion (85 μm nozzle) into 500 μL ice-cold 0.67*×* PBS + 2% FBS.

### 2.7 Single-cell library construction and sequencing

Sorted cells were pelleted at 300 × g for 5 min and counted by AO/PI (Invitrogen, cat. no. A49905). Cell density was adjusted to 1,500 cells/μL and loaded onto chips targeting 15,000 cells per animal using the 10x Genomics Chromium GEM-X Single Cell 5′ v3 Gene Expression platform. cDNA and library preparation followed manufacturer protocol CG000733 (RevA). Gene expression libraries were sequenced on an Illumina NextSeq2000 with a XLEAP-SBS P4 flow cell in a 28-10-10-90 configuration targeting 10,000 read pairs/cell; 2% PhiX was spiked in as a loading control.

### 2.8 Custom reference genome construction

For the initial 10x Genomics scRNA-seq preprocessing of *P. marinus*, we constructed a custom reference transcriptome from the Pmarinus 7.0 genome assembly and the matching Ensembl release 112 annotation. Because a primary assembly was not available, we used the top-level genome FASTA (.dna.toplevel.fa). We pre-filtered the annotation GTF to protein-coding genes with `cellranger mkgtf` (Cell Ranger v8.0.1; 10x Genomics) and built the reference with `cellranger mkref`. This initial counting reference was distinct from the later UTR-aware Ensembl+Evo2 reprocessing reference described above.

### 2.9 scRNA-seq data processing

Raw BCL files were demultiplexed into FASTQ format using `cellranger mkfastq` with one mismatch permitted during index demultiplexing. Reads were aligned, filtered, barcoded, and UMI-quantified against the custom *P. marinus* reference using `cellranger count` with default parameters.

### 2.10 Benchmarking on a degraded elephant shark genome

For robustness benchmarking, we also analyzed an elephant shark (*Callorhinchus milii* ) genome in which repetitive elements were aggressively hard-masked to simulate the fragmentation seen in lamprey. We first compared BRAKER2 and Evo2 *de novo* on the degraded genome at raw exon resolution, then evaluated a rescue benchmark in which structurally filtered Evo2 additions were merged onto a reduced Ensembl backbone after withholding 10% of protein-coding genes. Supplementary Methods gives the exact masking, withholding, and filtering settings.

## 3 Results

### 3.1 Workflow overview and robustness validation on a hard-masked elephant shark genome

A hybrid workflow carries predicted loci into rescue-space, pathway, transcription factor, and single-cell analyses (Fig. 1A). We stress-tested this workflow on a hard-masked elephant shark genome, where repeat masking removes much of the native exon signal and creates a degraded annotation substrate before we interpret lamprey-specific biology. Raw caller behavior and rescue-stage utility separate cleanly in this benchmark. At raw exon resolution, sensitivity denotes the fraction of Ensembl truth exons overlapped on the correct strand by at least one predicted exon, precision denotes the fraction of predicted exons that overlap a truth exon on the correct strand, and F1 is the harmonic mean of the two. Under that definition, BRAKER2 retained partial structural recovery on the hard-masked genome (sensitivity = 0.441, precision = 0.608, F1 = 0.511), whereas raw Evo2 exon calls remained noisy and low precision (sensitivity = 0.024, precision = 0.020, F1 = 0.022), so the raw exon benchmark favors BRAKER2 as the caller on the degraded genome (Fig. 1B). Evo2 contributes later as a filtered rescue layer rather than as a stronger raw exon caller. Withheld-gene rescue recall drops quickly as the structural threshold and minimum-length filters tighten (Fig. 1C). The full precision–recall tradeoff across the same settings, including the permissive regime (*S* ≥ 0.95, protein length ≥ 30 aa) that recovers more withheld genes but admits many weak additions, is shown separately (Supplementary Fig. S2). Under the strict benchmark setting (*S* ≥ 0.9745, protein length 100 ≥ aa), the merge contributed 413 non-overlapping additions and rescued 23 withheld genes, increasing full-truth gene coverage from 0.9002 to 0.9011 under severe masking (Supplementary Fig. S1). This elephant shark analysis serves as a lower-bound stress test rather than a direct estimate of lamprey performance. It does not support Evo2 as a stand-alone vertebrate annotator on hard-masked assemblies. Within this benchmark, BRAKER2 improves raw structure recovery, Evo2 contributes a filtered rescue layer, and neither replaces a transcript-boundary-aware backbone for scRNA quantification.

**Fig. 1.**
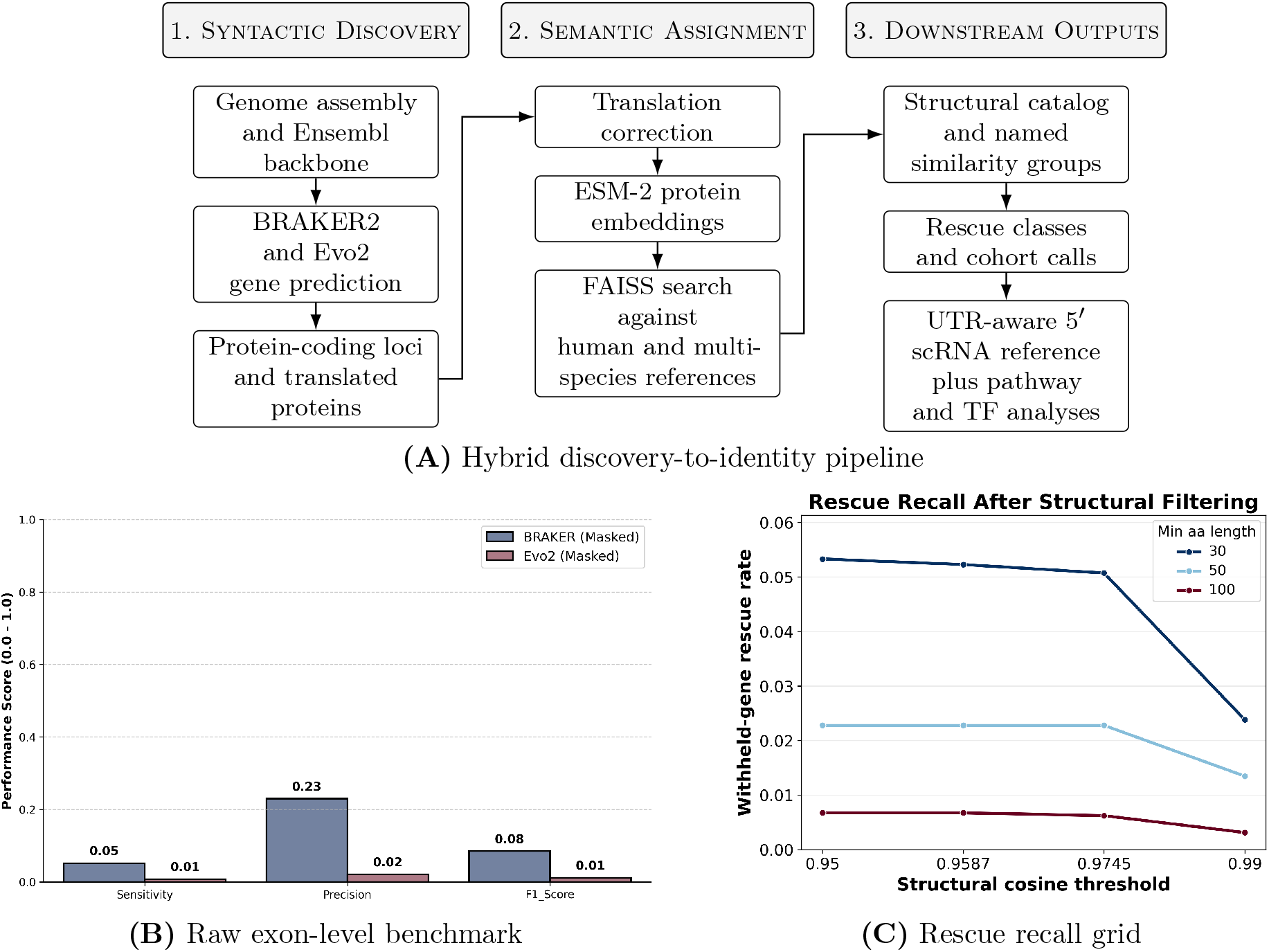
Workflow overview and early validation. (A) Schematic overview of the three-stage workflow: syntactic gene discovery, semantic cross-species similarity assignment in protein-embedding space, and downstream evolutionary stratification and validation. (B) Raw exon-level benchmark on a hard-masked elephant shark genome against Ensembl elephant shark annotations. BRAKER2 retains partial recovery on the degraded genome, whereas raw Evo2 exon calls remain highly permissive and low precision, so the raw exon benchmark favors BRAKER2. (C) Withheld-gene rescue recall across the elephant shark structural-threshold and minimum-length grid, showing that recall drops quickly as filtering tightens. The matched coverage summaries for the strict benchmark setting and the best merged setting are separated (Supplementary Fig. S1), and the full precision–recall tradeoff scatter is preserved separately (Supplementary Fig. S2).

### 3.2 Structural discovery and functional rescue

In lamprey, the workflow expands the structural catalog from the 10,415-locus Ensembl baseline to 31,286 catalog loci by adding 20,871 Evo2-derived loci absent from Ensembl (Fig. 2A). We treat that output as a structural catalog rather than a validated one-to-one gene count because structural support varies across the recovered set (Fig. 2B).

**Fig. 2.**
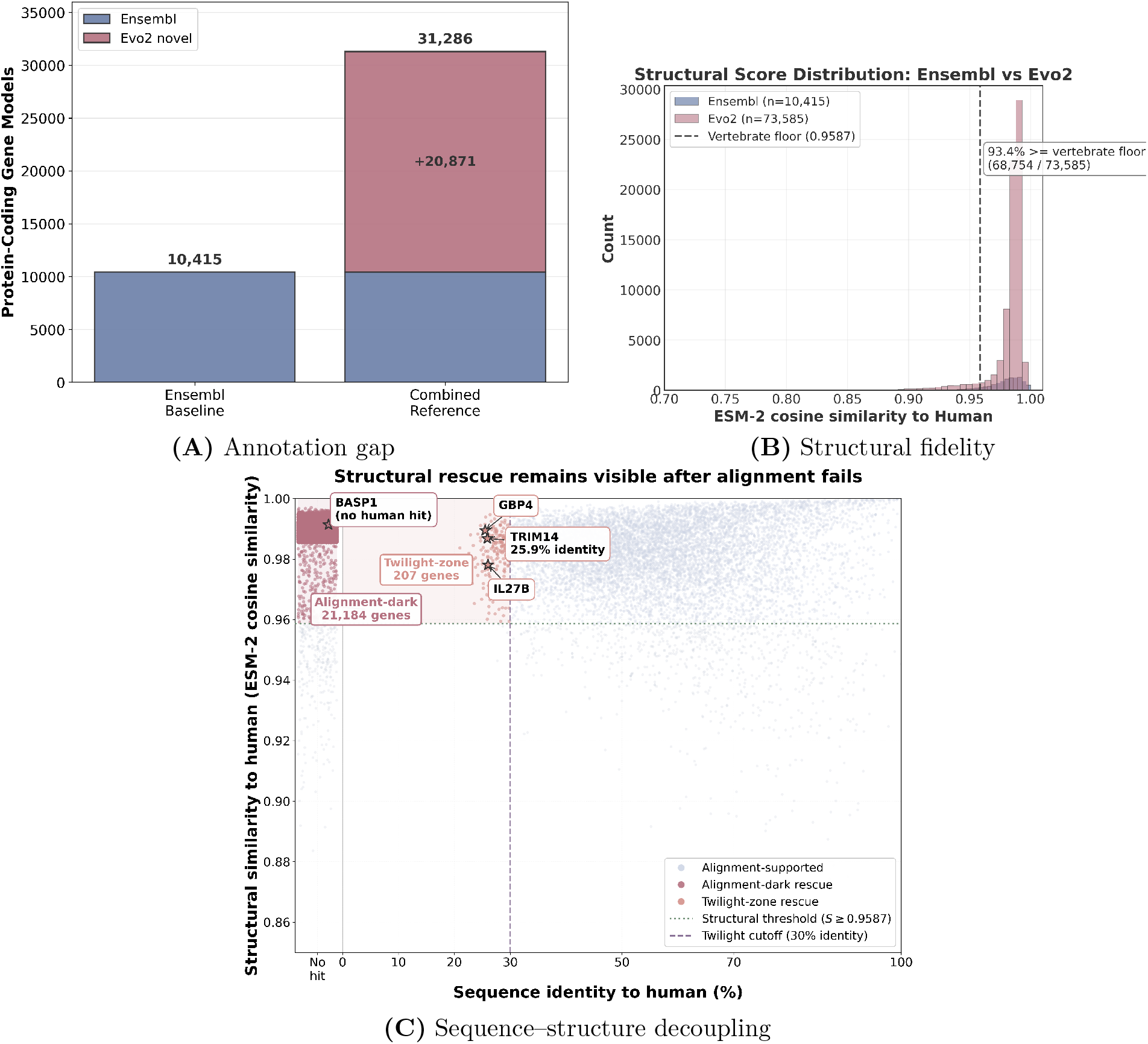
Catalog expansion and rescue-space structure. (A) Structural catalog composition: Ensembl baseline (10,415 genes) plus stacked Evo2 novel additions (20,871) yields a combined catalog of 31,286 loci. (B) Distribution-level view of structural fidelity across the recovered catalog. (C) Rescue-space scatter built from the current structural catalog. Alignment-dark catalog loci with no detectable human protein-sequence hit under our search settings occupy the left strip, while the measurable twilight-zone subset remains below 30% amino-acid identity but above the structural threshold. The labeled examples show both rescue modes directly, with enlarged callouts for *BASP1, TRIM14, IL27B*, and *GBP4*.

Much of that catalog signal comes from loci that remain well supported in embedding space even when protein-sequence alignment cannot recover them cleanly. Using the empirical vertebrate floor (*S* ≥ 0.9587), 21,391 catalog loci fall into this rescue space. Of those, 21,184 have no detectable human protein-sequence match under our search settings, and 207 retain only low-confidence human matches below 30% amino-acid identity. Those totals describe structurally supported catalog loci under a fixed human DIAMOND protein search, not one-to-one human-missing lamprey genes; duplication, fragmentation, and residual catalog noise can inflate locus counts relative to distinct conserved families. The two rescue classes map onto the two practical failure modes of protein-sequence alignment in divergent genomes. The anonymous Ensembl locus `ENSPMAG00000001780` has no detectable human protein-sequence match yet maps near the vertebrate neural regulator *BASP1* in embedding space, consistent with a candidate cross-species family relationship. `ENSPMAG00000004750` retains only 25.9% amino-acid identity to human but still maps near *TRIM14*. The no-hit strip and the measurable twilight-zone cloud show those two regimes directly (Fig. 2C). Supplementary Tables S2A–S2B list representative loci from both groups. Additional immune-linked similarity assignments in the twilight-zone regime include loci mapping near *IL27B* (*S* = 0.9781, 26.0% amino-acid identity) and *GBP4* (*S* = 0.9895, 25.5% amino-acid identity). Structural embeddings therefore recover candidate cross-species family relationships across deep divergence both when protein-sequence alignment returns a weak hit and when it fails outright.

### 3.3 Vertebrate-biased structural cohort

Evolutionary stratification partitions the named structural catalog into bilaterian core, a vertebrate-biased cohort, and lamprey-specific cohorts. Under the current phylogenetic classification, the named catalog contains 6,143 unique cross-species similarity groups represented by 84,000 annotated loci, and 1,739 of those groups represented by 26,647 loci fall into the vertebrate-biased cohort under this structural classification. Projecting those groups into phylogenetic structural space separates bilaterian core genes from vertebrate-biased loci in an evolutionary map (Fig. 3A,B).

**Fig. 3.**
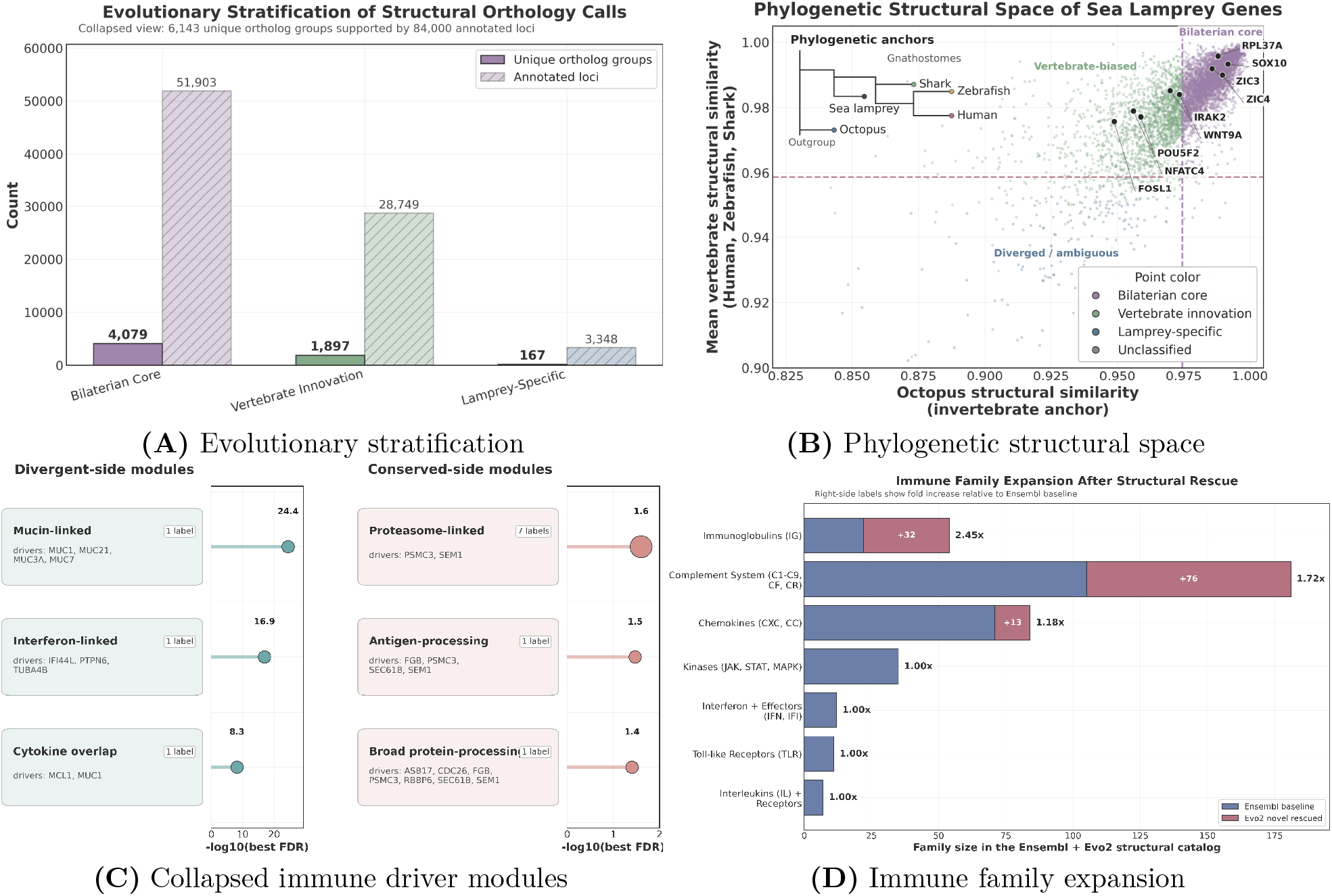
Vertebrate-biased structural cohort and its immune-functional structure. (A) Stratification of named structural similarity calls at group and annotated-locus levels. (B) Phylogenetic structural space of sea lamprey cross-species similarity groups, positioned by mean vertebrate similarity versus Octopus similarity and colored by inferred evolutionary class. (C) Immune Reactome terms collapsed by shared lamprey driver genes; cards name the shared drivers, badges report the number of terms per module, and terminal markers show the best FDR. (D) Immune family expansion after high-confidence structural rescue relative to the Ensembl baseline.

This vertebrate-biased set is enriched for immune-linked similarity assignments, including interleukin-like and chemokine-like families. The Reactome analysis shows the same pattern: nuclear receptor transcription, Cdc42 GTPase signaling, and gap junction assembly stay conserved, while sensory- and translation-linked categories show the strongest outgroup-biased losses (Supplementary Fig. S3). Overlapping human Reactome terms were collapsed by shared mapped lamprey driver genes, so one repeated two-gene signal does not appear as several pathways (Fig. 3C).

After collapsing overlapping terms, six modules remain. On the conserved side, seven terms collapse onto a proteasome-linked module anchored by *PSMC3* and *SEM1*, and two smaller modules add *SEC61B* with broader protein-processing hits. On the divergent side, three compact modules carry the signal, driven by mucin-like hits (*MUC1, MUC21, MUC3A, MUC7* ), *IFI44L*/*PTPN6* /*TUBA4B*, and *MCL1* /*MUC1*. Lamprey lacks the canonical mammalian IgE and Fc*ε* receptor genes, so those Reactome terms arise from associated pathway genes such as *PSMC3, SEM1*, and *SEC61B* rather than from those canonical mammalian genes themselves. We therefore interpret these similarity-mapped modules as proteostasis, antigen-processing, and cytokine-linked programs rather than as one-to-one mammalian IgE, Fc*ε* receptor, immunoglobulin, or MHC pathways. The same pattern appears at the family level, where a small set of immune classes expands after structural rescue instead of the catalog inflating evenly (Fig. 3D). Together, the module and family-expansion panels identify the immune signal before the phylogenetic comparison traces it across species (Fig. 3C,D).

### 3.4 Phylogenetic pathway trajectories place those immune modules in context

The immune signal first collapses into a small set of shared driver modules, then those same modules can be traced across the outgroup-to-vertebrate series (Fig. 3C; Fig. 4). Ancient cellular pathways such as cell cycle and RNA metabolism stay uniformly high across the tree, consistent with deep bilaterian conservation (Fig. 4A). Similarity-mapped modules labeled by human Reactome terms such as TLR3-, interferon-, and CD28-related signaling show vertebrate-biased structural shifts, with higher similarity to elephant shark, zebrafish, and human than to octopus (Fig. 4B). Receptor and sensory pathways such as tachykinin, P2Y, and LGI-ADAM show the strongest outgroup drop (Supplementary Fig. S3B). Immune labels accumulate differently across the two directions: conserved labels cluster around receptor-like, TCR, and antigen-presentation terms, whereas divergent labels spread across interferon-, interleukin-, neutrophil-, and Dectin-linked terms (Fig. 4C). Taken together, the module, family-expansion, and trajectory panels show that the strongest remodeling sits in selected immune and receptor programs rather than across the pathway landscape as a whole (Fig. 3C,D; Fig. 4).

**Fig. 4.**
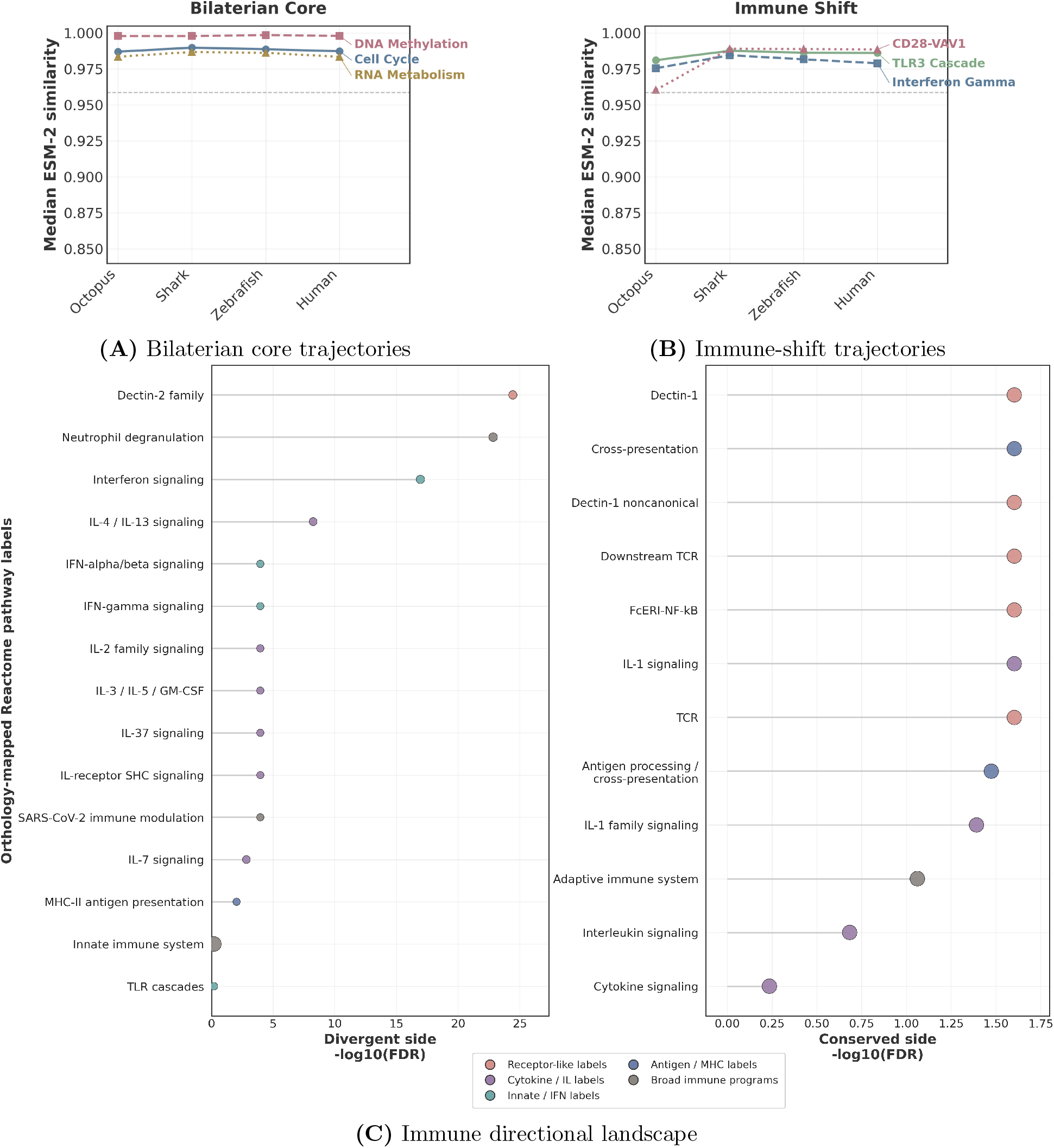
Phylogenetic structural profiles expose branch-specific remodeling. (A) Bilaterian-core trajectories remain uniformly high across the outgroup-to-vertebrate series. (B) Representative immune pathways show vertebrate-biased shifts. (C) Directional landscape of human-mapped immune Reactome terms, split by enrichment direction and plotted by pathway-level significance; dot size indicates gene-set size and color indicates immune module family. Conserved-side terms cluster around receptor-like, TCR, and antigen-presentation categories, whereas divergent-side terms span interferon-, interleukin-, neutrophil-, and Dectin-linked categories. Terms are human Reactome names transferred by similarity mapping rather than literal one-to-one mammalian pathway claims.

### 3.5 Transcription factor architecture is broadly conserved with targeted vertebrate shifts

To ask whether the same branch structure extended to regulatory genes, we analyzed the transcription factor repertoire. Using the curated Lambert/Weirauch TF census, we identified 365 transcription factor groups spanning 820 annotated lamprey loci in the Ensembl+Evo2 consensus set. Most of the repertoire remained close to the bilaterian core. Large TF superfamilies such as C2H2 zinc fingers, bHLH factors, and most ETS factors showed conservation rather than wholesale rewiring.

The measurable branch shifts were family-specific and modest. Restricting attention to families represented by multiple groups, Grainyhead showed the largest vertebrate-biased shift (median Δ = 0.017; 3/4 groups classified within the vertebrate-biased cohort), while Homeodomain/POU, bZIP, bHLH, and ETS families showed smaller vertebrate enrichments (Supplementary Methods S7). At the single-gene level, the highest-ranked vertebrate-biased TF names in this census included developmental and immune-linked regulators such as *ONECUT2, SATB2, UBP1, FOSL1, POU5F2*, and *NFATC4* (Supplementary Table S3). The lamprey regulatory toolkit remains largely ancient, and the branch-specific shifts stay confined to a limited subset of developmental and immune-linked factors rather than extending to genome-wide TF turnover.

### 3.6 Single-cell reanalysis refines annotation of adaptive and iron-linked states in the lamprey immune compartment

We re-analyzed the reclustered immune subset from the whole-blood atlas with the UTR-aware single-cell RNA-seq reference to test whether the expanded annotation changed biological interpretation. Prior immune-cluster labels together with immune-marker expression, including *ptprc*/CD45 (`ENSPMAG00000003196`), defined the subset. In lamprey, adaptive immune states are commonly described by their variable lymphocyte receptor (VLR) programs, so we use VLRA^+^ and VLRB^+^ to denote clusters enriched for transcripts associated with variable lymphocyte receptor A or B programs, respectively, rather than direct receptor measurements (Pancer et al. 2004; Guo et al. 2009). Because additional VLR lineages exist, these labels do not exclude other receptor-defined states from the same clusters. These are cluster-level marker programs rather than one-to-one mammalian cell-type assignments. Panel A in Fig. 5 shows the consensus atlas view of that same reprocessed immune subset, using the hybrid Ensembl+Evo2 reference and the shared C-cluster labels that anchor the comparisons in panels B and C. The Ensembl+Evo2 reference preserved the overall immune manifold and expanded the marker space available for annotation. Cell recovery stayed similar between the Ensembl+Evo2 and Ensembl references after preprocessing (33,277 versus 33,403 globally; 2,383 versus 2,385 in the immune subset), while gene recovery after QC increased from 7,484 genes to 8,455 genes. At the raw Leiden level, the Ensembl-only reference resolved five immune partitions, whereas the hybrid reference condensed the same immune compartment into four and yielded 600 rather than 720 unique top-50 DEG genes, consistent with cluster consolidation rather than loss of information. The hybrid atlas still contains a 92-cell iron-associated VLR-linked cluster, but the updated annotation no longer splits that population into a separate fifth Leiden partition. Overall, the hybrid reference changed the named-feature space and simplified the cluster structure more than it changed cell recovery.

**Fig. 5.**
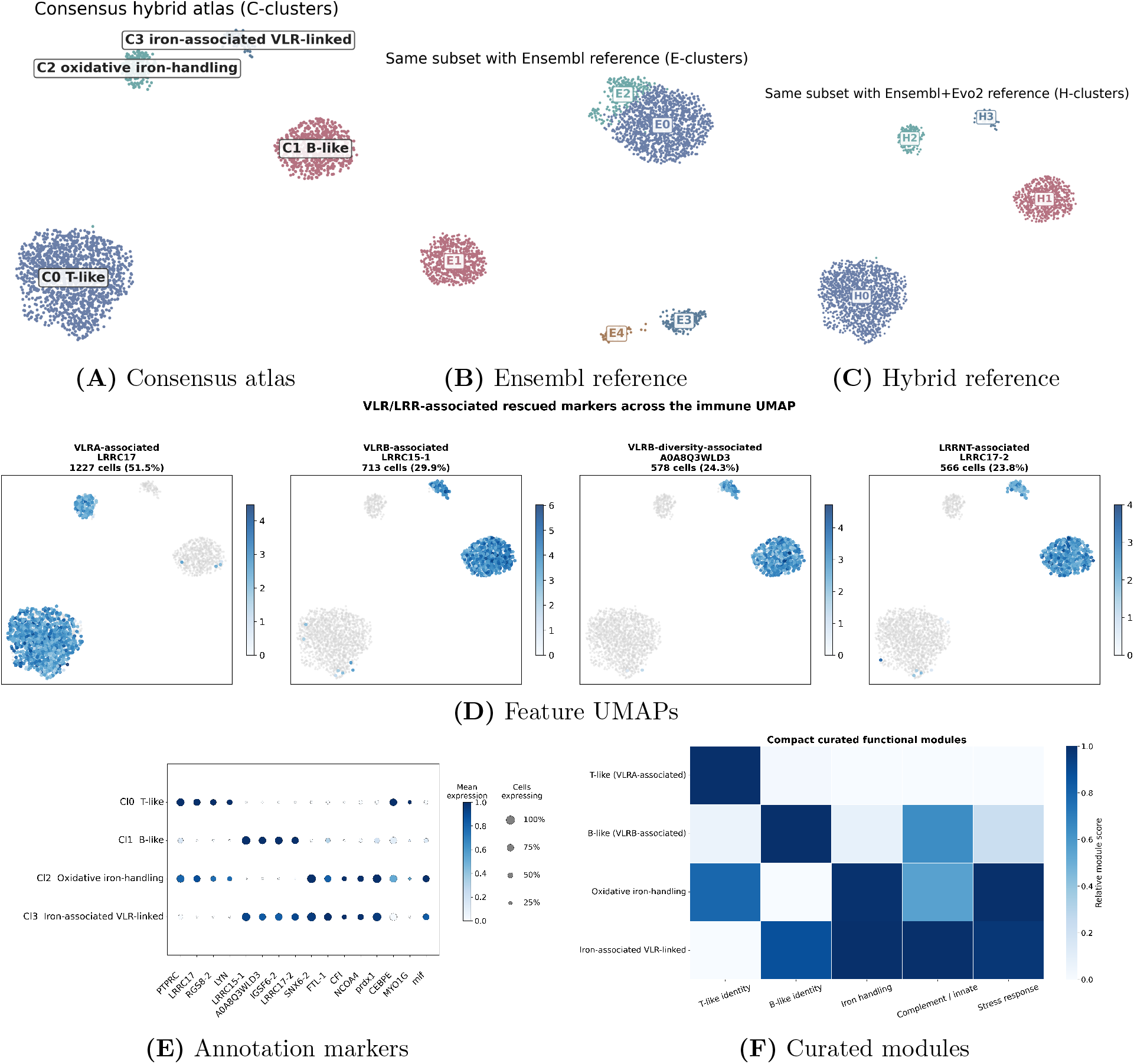
scRNA-seq validation of the immune compartment. (A) Consensus atlas view of the same reprocessed whole-blood immune subset used throughout this figure, shown after counting with the hybrid Ensembl+Evo2 reference and annotated with the shared C-cluster labels used for cross-panel comparison. (B) The same immune subset counted with the Ensembl-only reference. (C) The same immune subset counted with the Ensembl+Evo2 hybrid reference. Together, panels A–C show that the hybrid reference preserves the overall immune manifold while consolidating the five-way Ensembl-only Leiden partition into four hybrid partitions and preserving marker-based annotation. In these UMAP panels, C labels mark the consensus atlas clusters, E labels mark Ensembl-only cluster IDs, and H labels mark hybrid-reference cluster IDs. (D) Feature UMAPs of VLR- and LRR-associated markers that localize the VLRA^+^- and VLRB^+^-linked programs. (E) Dot plot of defining immune markers used for cluster annotation, including structurally supported markers that distinguish the oxidative and iron-associated states. (F) Curated module-score heatmap summarizing T-like identity, B-like identity, iron handling, complement/innate activity, and stress response across the four immune states; module members are listed in Supplementary Table S4.

With that terminology, the hybrid reference supports annotation of four transcriptionally defined immune populations. These include a dominant VLRA^+^ T-like cluster, a VLRB^+^ B-like cluster, an oxidative iron-handling state enriched for ferritin-linked and redox-response markers, and an iron-associated VLR-linked state. The latter retains ferritin-linked and redox-linked programs while also carrying structurally supported B-like markers such as *LRRC15, IGSF6*, and *CFI*. Both smaller states sit next to the VLR-associated adaptive compartment but do not map cleanly onto one-to-one mammalian analogs, so we describe them conservatively rather than assigning a specific effector behavior in vivo.

Structural rescue contributes materially to this annotation. Across the top 25 markers per cluster, ESM-2-rescued names accounted for 15/25, 10/25, 14/25, and 15/25 markers in the VLRA^+^-associated T-like, VLRB^+^-associated B-like, oxidative iron-handling, and iron-associated VLR-linked clusters, respectively; including anonymous structurally supported loci increased those fractions to 17/25, 10/25, 14/25, and 15/25.

The rescued-marker burden helps explain those cluster annotations. Feature UMAPs of rescued variable lymphocyte receptor- and leucine-rich-repeat-associated markers place the expected VLR-associated subspaces (Fig. 5D), and the annotation-marker dot plot shows that structurally supported marker names help distinguish the VLRA^+^ and VLRB^+^ programs from the oxidative and iron-associated states (Fig. 5E). Adaptive-state module scores remain highest in the canonical T-like and B-like clusters, ferritin-linked and oxidative scores peak in the oxidative iron-handling cluster, and the iron-associated VLR-linked cluster combines B-like, iron-handling, complement/innate, and stress-linked signatures (Fig. 5F). Supplementary Table S4 lists the gene sets used to score those curated modules. Cluster-level provenance follows the same split between baseline Ensembl names, rescued calls, anonymous Ensembl IDs, and novel Evo2 loci (Supplementary Fig. S4). Suffixes such as ‘LRRC17-1’ denote separate lamprey loci that share the same broad cross-species family label rather than transcript isoforms. Doublet analysis (UMI ratios 1.07–1.11×; per-cell co-expression remained weak, |*r*| ≤ 0.22; Supplementary Fig. S5) argues against technical admixture and is consistent with these clusters representing mixed-function immune states in the lamprey blood compartment. We note that multiple T-like VLR lineages exist in lamprey, including VLRC and additional recently described VLR classes (Guo et al. 2009; Das et al. 2023, 2025); therefore, the VLRA^+^-associated cluster identified here likely reflects a broader T-like transcriptional state rather than a one-to-one mapping to a single receptor-defined lineage.

## 4 Discussion

Highly divergent genomes still create a naming problem after structure prediction. Gene-structure predictors can recover exon–intron boundaries and coding coordinates, but cross-species similarity mapping remains difficult once pairwise comparisons among translated proteins lose signal. Our framework tackles that post-discovery step by separating gene-model discovery from cross-species similarity mapping. Across 73,585 Evo2-derived translated protein models, 73,485 received high- or medium-confidence cross-species similarity assignments, including 35,395 high-confidence calls. After collapsing loci that shared the same resolved cross-species similarity group, those assignments add 20,871 Evo2 catalog loci beyond Ensembl and expand the lamprey structural catalog to 31,286 loci while leaving room for a stricter UTR-aware subset for scRNA quantification.

Most of that gain comes from loci that embeddings rescue after pairwise protein-sequence alignment weakens or disappears. That pattern suggests that deep cyclostome proteins often retain usable structural similarity after amino-acid similarity drops below the range where standard transfer pipelines work. The immune-linked similarity assignments in the vertebrate-biased structural cohort point in the same direction. The catalog contains cytokine-like, chemokine-like, and antigen-processing similarity assignments that remain difficult to recover with pairwise protein-sequence alignment alone. The transcription factor analysis reinforces that pattern. Regulatory architecture stays mostly ancient, and the branch-specific shifts affect a limited subset of developmental and immune-linked families rather than genome-wide TF turnover. We emphasize that the large rescue-space in lamprey reflects an annotation-space expansion in a deeply divergent and repetitive genome, and should not be interpreted as a direct estimate of distinct conserved genes.

Concurrent with this work, ANNEVO showed that evidence-free deep models can achieve *ab initio* gene-structure prediction across broad phylogenetic ranges. That advance changes the discovery side of the problem, not the naming side. BRAKER2 uses external evidence to sharpen exon–intron boundaries and transcript coordinates. ANNEVO predicts those boundaries from genomic sequence alone. Our framework starts after that step and asks a different question: once a model proposes a protein, can we place it in phylogenetic structural space, derive a plausible cross-species similarity mapping, quantify rescue beyond pairwise protein-sequence alignment, and show that the recovered annotation improves downstream analyses such as scRNA-seq interpretation? Genomic-sequence-only structure prediction still leaves those semantic and downstream questions open. The separation between discovery and the similarity-assignment layer should therefore remain useful even as stronger discovery engines appear.

The single-cell analysis supports the same conclusion. CDS-oriented *ab initio* models, whether generated by Evo2, BRAKER2, or newer deep predictors, cannot serve as scRNA-seq references without transcript-boundary-aware backbones. In 10x 5′ data, reads concentrate near transcript starts, first exons, and 5′ untranslated regions, which CDS-only models often omit. In the lamprey immune compartment, the hybrid reference supports annotation of two iron-linked states adjacent to VLRA^+^-associated and VLRB^+^-associated adaptive clusters. Here, “VLRA^+^-associated” and “VLRB^+^-associated” denote clusters enriched for transcripts mapping to those receptor programs rather than direct measurement of receptor expression at the protein level. Because multiple VLR genes exist in lamprey beyond VLRA^+^ and VLRB^+^, we interpret these clusters as broader T-like and B-like transcriptional states rather than exhaustive receptor-defined populations. These data place them on an iron-linked continuum adjacent to the VLR-associated adaptive compartment, but they do not yet resolve whether they define a distinct cyclostome immune program or part of a broader erythroid or iron-recycling spectrum. More broadly, the hybrid reference assigns names to previously anonymous features and yields testable evolutionary and cellular hypotheses.

## Limitations

The Evo2 structural catalog still contains noise from pseudogenes, non-coding RNAs with coding-like syntax, and imprecise start codon prediction. Across the Evo2 semantic input set, 73,585 translated protein models reached the assignment stage; 35,395 received high-confidence assignments, 38,090 received medium-confidence assignments, and 100 remained low-confidence or unassigned. Those model-level totals are much larger than the final deduplicated gene catalog and should be read as semantic inputs rather than distinct gene counts. The same caution applies to the rescue-space summary. The 21,391-locus total is a catalog-level count under fixed human search settings, not a one-to-one estimate of biologically distinct human-absent lamprey genes. The elephant shark masking benchmark establishes a lower bound on recovery when sequence information is degraded or missing, but it does not fully reproduce the evolutionary setting in which sequence remains present yet pairwise alignment no longer resolves relationships cleanly across lineages. Proteomics and long-read RNA-seq will be needed to validate the full filtered set. Direct benchmarks against newer evidence-free predictors are the next test, especially if we want to measure how much value the semantic assignment layer adds when paired with stronger structural predictors. We expect the same framework to transfer to other divergent lineages where alignment-based methods have stalled.

## Future directions

Immediate extensions of this framework include benchmarking in more closely related but incompletely annotated vertebrate genomes, incorporating long-read transcriptomic and proteomic evidence to refine structural calls, and testing richer similarity summaries that move beyond a single best structural match. A phylogenetically closer benchmark such as shark would be especially useful because it would test cross-species relationship recovery in a regime where sequence remains present but standard alignment-based transfer has become unreliable. More broadly, the same rescue-stage design should be useful in other deeply divergent systems where transcript-guided backbones exist but pairwise protein-sequence alignment alone leaves substantial fractions of the coding repertoire unnamed. When pairwise comparisons among translated proteins stall in a divergent genome, structural discovery plus similarity assignment can still recover biologically informative annotations and expose questions that are otherwise difficult to ask.

## Acknowledgments

We thank the Dartmouth Discovery cluster for GPU and HPC support. We thank the U.S. Geological Survey Hammond Bay Biological Station for assistance with *P. marinus* procurement, and Grigoriy Losyev from the Brigham and Women’s Hospital (BWH) Flow Core for assistance with flow cytometry and fluorescence-activated cell sorting (FACS). We acknowledge the animals used in this study and recognize their essential contribution to the advancement of scientific knowledge. All animal procedures were conducted in accordance with institutional and ethical guidelines.

## Declarations

### Author contributions

T.B.L., S.K.C., D.R.W., and H.R.F. conceived and designed the research. G.A.P., J.W.C., M.H., and M.K. performed experiments. T.B.L. and S.S. analyzed the data. T.B.L., S.K.C., G.A.P., J.W.C., S.S., D.R.W., and H.R.F. interpreted the results. T.B.L. drafted the manuscript. T.B.L., S.K.C., G.A.P., J.W.C., S.S., D.R.W., and H.R.F. edited and revised the manuscript and approved the final version.

### Ethics approval and consent to participate

All animal procedures were approved by the Brigham and Women’s Hospital Institutional Animal Care and Use Committee under protocol 2023N000131.

### Data and code availability

The sea lamprey scRNA-seq data are deposited under BioProject `PRJNA1459322`. NCBI lists the current submission as `SUB16137332`; the data will be released upon publication. The code and manuscript source for this study are available at https://github.com/tobylanser/bioFM annotation.

