## Supplemental methods and materials for "Biological foundation models illuminate annotation blind spots in evolutionarily divergent genomes"

**Abstract**

This file contains the supplementary methods for the lamprey annotation study, including model configuration, cross-species similarity assignment thresholds, rescue-space classification, pathway analysis, transcription factor conservation analysis, and elephant shark validation benchmark details.

### Supplementary Methods

#### S1. Model Architectures and Parameters

##### S1.1 Evo2: Genomic Language Model

We employed Evo2, a genomic foundation model trained across all domains of life (Brixi et al. 2026).

| Parameter | Value |
|---|---|
| Model variant | evo2-7b |
| Context window | 131,072 bp |
| Tokenization | Single nucleotide (1 bp) |
| Attention mechanism | Rotary position embedding (RoPE) |
| Inference precision | bfloat16 |
| GPU requirements | 1× L40S or H100 (48–80GB) |

**Table 1** Evo2 inference parameters.

***Streaming deployment.***

To remain within storage limits, we used a streaming inference architecture based on the official Evo2 exon-classifier workflow. Genomic windows (8,192 bp, 100 bp overlap) were processed in VRAM, intermediate Evo2 block-26 embeddings were extracted on both strands, and the forward/reverse embeddings for each genomic position were concatenated and scored with the lightweight `schmojo/evo2-exon-classifier`. We retained exon calls and their summary scores rather than storing whole-genome embedding dumps.

***Dual-strand inference and boundary refinement.***

We ran inference on both the forward and reverse-complement strands for every locus to reduce strand ambiguity. Exonic positions were called at classifier probability $\geq 0.5$, merged across gaps $< 20$ bp on the same strand, and then expanded by a heuristic 25 bp upstream / 50 bp downstream boundary adjustment for downstream locus recovery. These settings are post-processing heuristics, not learned Evo2 model parameters.

***Post-processing.***

Raw Evo2 predictions occasionally merged adjacent genes into megagenes (spanning $> 1$ Mb). We split gene models at intergenic gaps $> 500$ bp using `utils/format_evo2_gtf.py`, producing biologically plausible loci while preserving exon structure within each fragment.

#### S1.2 ESM-2: Protein Language Model

We used ESM-2 (`esm2_t33_650M_UR50D`), a 650M-parameter encoder-only transformer.

| Parameter | Value |
|---|---|
| Model variant | esm2_t33_650M_UR50D |
| Hidden dimension | 1,280 |
| Attention heads | 20 |
| Layers | 33 |
| Training data | UniRef50 (65M sequences) |
| Embedding extraction | Layer 33, mean pooling |
| Max sequence length | 1,024 residues |

**Table 2** ESM-2 architecture and inference parameters.

***Translation and preprocessing.***

Predicted gene models were translated using a robust multi-frame ORF selection strategy implemented with Biopython sequence utilities and preferentially retaining long ORFs with canonical start codons (Cock et al. 2009). Sequences containing internal stop codons (`*`) or ambiguous residues were cleaned before embedding. For ESM-2 inference, proteins longer than the effective 1,024-token limit were truncated to 1,022 residues before mean pooling of the final-layer representations; the workflow used here does not yet implement sliding-window representation averaging for long proteins.

#### S1.3 FAISS: Vector Similarity Search

Cross-species similarity searches were performed with FAISS using cosine similarity (via L2 normalization) and exact inner-product indexing.

| Parameter | Value |
|---|---|
| Index type | IndexFlatIP (exact search) |
| Distance metric | Cosine similarity (L2-normalized IP) |
| Vector dimension | 1,280 |
| Reference proteomes | Human, Zebrafish, Elephant Shark, Octopus |
| Query vectors (Lamprey) | 73,585 (Evo2 proteins) |

**Table 3** FAISS search configuration.

##### S1.4 Reference proteomes and identifier namespaces

The structural search used four external reference proteomes: Human, Zebrafish, Elephant Shark, and Octopus. We used Human UniProt reference proteome `UP000005640` (GRCh38.p14), Zebrafish reference proteome `UP000000437` together with the Ensembl GRCz11 genome/annotation backbone used for the stress test (release 113), Elephant Shark UniProt reference proteome `UP000314986` (assembly `Callorhinchus_milii-6.1.3`), and Octopus (*Octopus sinensis*) UniProt reference proteome `UP000515154` (assembly `O.sinensis_v1.0`). The lamprey baseline annotation was anchored to Ensembl release 110 on assembly `Pmarinus_7.0` (`GCA_000466065.2`). In the comparative design, Elephant Shark (*Callorhinchus milii*) served as the cartilaginous-fish gnathostome anchor. This choice was motivated by prior comparative-genomics work describing the elephant shark genome as unusually slowly evolving and therefore especially informative for reconstructing early jawed-vertebrate biology (Venkatesh et al. 2014).

Lamprey features kept Ensembl stable identifiers throughout the workflow, including `ENSPMAG` gene IDs, `ENSPMAT` transcript IDs, and `ENSPMAP` protein IDs. The consensus tables separate display names from provenance. The top-hit columns for Human, Zebrafish, Elephant Shark, and Octopus retain the native accession systems of those reference proteomes, `source` records which vertebrate reference supplied the best-scoring hit, `best_source_hit` records the literal winning sequence label, and `final_name` provides the display symbol once the locus exceeds the vertebrate similarity floor. That separation keeps pathway analyses and scRNA dotplots readable without hiding where each name came from. When multiple lamprey loci collapse onto the same broad cross-species similarity family, suffixes such as `-1` mark separate loci; in this workflow they do not denote transcript isoforms.

#### S2. Consensus Cross-Species Similarity Assignment

For each Evo2 protein embedding, we performed FAISS search against each reference proteome and recorded the top-1 hit and similarity score. Consensus identities were then assigned from the best vertebrate-supported match rather than from a fixed source order:

- **High-confidence vertebrate similarity assignment**: if the best score among Human, Zebrafish, and Elephant Shark satisfied $S \geq 0.9587$, that vertebrate source was taken as the primary structural match.
- **Ambiguity downgrade**: if the best vertebrate hit exceeded the threshold but the runner-up vertebrate score fell within 0.005 cosine-similarity units, the locus was downgraded from `High` to `Medium` to indicate that the cross-species assignment was structurally close but not decisively separable.
- **Lamprey self-hit fallback**: if no vertebrate hit cleared the vertebrate floor and an Ensembl lamprey self-hit existed with $S_{Lamprey} \geq 0.98$, the Ensembl symbol was retained.
- **Medium-confidence fallback**: if no stronger rule applied but the Human score satisfied $S_{Human} \geq 0.85$, the locus was retained as a medium-confidence similarity assignment.

We write these annotations to `evo2_consensus_annotation.csv` with the top hits and scores for all reference species, together with the winning vertebrate source, the literal winning accession, and the final display name used in downstream analyses.

#### S3. Threshold Calibration

##### S3.1 Empirical null distributions

We estimated null similarity distributions using random non-homologous protein pairs (Human vs Octopus; Human vs Lamprey) to derive percentile-based thresholds. The vertebrate floor was set at the 99th percentile of the Human-vs-Lamprey empirical null ($S = 0.9587$), and the bilaterian floor at the 99th percentile of the Human-vs-Octopus empirical null ($S = 0.9745$).

##### S3.2 Validation against known orthologs

We checked these percentile-derived cutoffs against curated vertebrate ortholog sets to confirm that the vertebrate floor retained most obvious vertebrate matches while remaining conservative against random background. We used these ortholog-set comparisons as validation rather than as the primary optimization criterion; the thresholds reported here come from the empirical nulls above.

##### S3.3 Full rescue-space classification

To distinguish strict twilight-zone cases from loci that are completely invisible to alignment, we constructed a canonical alignment–structure classification on the 31,286-entry manuscript structural catalog (10,415 Ensembl loci + 20,871 novel Evo2 loci). The structural catalog followed the main manuscript definition. It included all Ensembl consensus loci plus the top 20,871 non-redundant high-confidence Evo2 loci after removing names already present in Ensembl. For sequence search, we used the exact same catalog universe, with one longest-protein representative per Ensembl gene and the matched Evo2 protein for each retained novel locus.

Human sequence recoverability was then evaluated with a single consistent DIAMOND search (`diamond` v2.1.9; local project install) against the Human UniProt reference proteome. Query sequences were written to a combined FASTA built from the manuscript structural catalog, and DIAMOND was run in `blastp` mode with `--sensitive`, `--max-target-seqs 1`, and `--evalue 1e-3`. Output fields were `qseqid`, `sseqid`, `pident`, alignment coordinates, `evalue`, and `bitscore`. For each lamprey locus, only the top-scoring human hit was retained.

We then partitioned loci using two thresholds:

- **Strong structural support**: Human ESM-2 cosine similarity $S \geq 0.9587$.
- **Twilight-zone sequence regime**: best-hit human sequence identity $< 30\%$.

Loci were classified into four mutually exclusive groups:

- **Alignment-supported**: measurable human sequence hit with $\geq 30\%$ identity and $S \geq 0.9587$.
- **Twilight-zone rescue**: measurable human sequence hit with $< 30\%$ identity and $S \geq 0.9587$.
- **Alignment-dark rescue**: no measurable human sequence hit under our search settings, but $S \geq 0.9587$.
- **Low structure / unassigned**: all remaining loci.

Under this classification, the full rescue space is the union of Twilight-zone rescue and Alignment-dark rescue. On the 31,286-entry structural catalog, that yielded 21,391 rescued loci, with 207 strict twilight-zone cases and 21,184 alignment-dark loci. These are catalog-level locus counts under a fixed human DIAMOND search, not experimentally validated or one-to-one human-absent lamprey genes. Duplication, fragmentation, and multiple lamprey loci mapping to the same vertebrate family can all inflate locus totals relative to distinct conserved genes. We wrote the per-locus classifications to `RESULTS/tables/full_rescue_space_classification.csv`, with summary counts in `RESULTS/tables/full_rescue_space_summary.csv`.

For interpretability in the supplement, representative rescued loci were split into two tables rather than pooled into a single list. Supplementary Table S2A contains the top twilight-zone genes with measurable sequence identity below 30%, and Supplementary Table S2B contains the top alignment-dark genes with no measurable human hit under the same search settings.

#### S4. BRAKER2 Configuration

BRAKER2 was run in protein-only mode with softmasked genome input:

```
braker.pl \
  --genome=genome_softmasked.fa \
  --prot_seq=proteins.clean.fasta \
  --softmasking \
  --species=lamprey_braker \
  --cores=48 \
  --gff3
```

The protein database included Human (UniProt), Zebrafish (UniProt), Elephant Shark (combined UniProt and Ensembl-derived proteome), and OrthoDB v11 vertebrate proteins.

#### S5. scRNA-seq experimental methods, preprocessing, and reference construction

##### S5.1 Animal source, husbandry, and ethics

*Petromyzon marinus* larvae sea lamprey (9.5–10 cm long; 0.85–1.0 g) were obtained from the U.S. Geological Survey Hammond Bay Biological Station in Michigan. We maintained animals in a temperature-controlled tank with aerated filtered water at 17–21°C and pH 7.5–8.3. All animal studies were approved by the Brigham and Women's Hospital Institutional Animal Care and Use Committee under protocol 2023N000131.

##### S5.2 Cell preparation and sorting

We euthanized larval *P. marinus* in tricaine methanesulfonate buffered in Tris-HCl (pH 8.0) until they no longer responded to tail pinch. We collected blood from the tail vein and then decapitated the animals. Collected blood was centrifuged at $300\times g$ for 5 min and resuspended in 1 mL RBC lysis buffer (Invitrogen, cat. no. 00-4333-57). After a 5 min incubation at room temperature, cells were pelleted, washed with 1 mL $0.67\times$ PBS + 1% FBS (L-FACS buffer), passed through a 40 $\mu$m mesh filter, and pelleted again. Cells were then stained for 20 min on ice with 2.5 $\mu$g/mL 4′,6-diamidino-2-phenylindole (DAPI) in L-FACS buffer. We added 1 mL L-FACS buffer, pelleted the cells, resuspended them in 200 $\mu$L L-FACS buffer, and filtered them again through a 40 $\mu$m mesh. DAPI-negative live cells were sorted on a FACSAria Phusion with an 85 $\mu$m nozzle into tubes containing 500 $\mu$L ice-cold $0.67\times$ PBS + 2% FBS.

##### S5.3 Single-cell library construction and sequencing

Sorted cells were pelleted at $300\times g$ for 5 min, and an aliquot was counted with AO/PI (Invitrogen, cat. no. A49905). Cell density was adjusted to 1,500 cells/$\mu$L, and cells were loaded on Chromium GEM-X chips with a target recovery of 15,000 cells per animal using the 10x Genomics Chromium GEM-X Single Cell 5′ v3 Gene Expression platform. cDNA and library preparation followed the manufacturer's user guide (CG000733, Rev A). Gene-expression libraries were sequenced on an Illumina NextSeq 2000 with an XLEAP-SBS P4 flow cell using a 28-10-10-90 read configuration and a target depth of 10,000 read pairs per cell. We added 2% PhiX as a loading control.

##### S5.4 Custom reference genome construction

For the initial 10x Genomics single-cell RNA-seq preprocessing of sea lamprey, we generated a custom reference transcriptome from the *P. marinus* assembly `Pmarinus_7.0` and the matching Ensembl release 112 annotation. Because a primary assembly was not available for this species, we used the top-level genome FASTA (`.dna.toplevel.fa`). Before reference generation, we filtered the annotation GTF to retain only protein-coding genes with `cellranger mkgtf` (Cell Ranger v8.0.1; 10x Genomics). We then built the custom reference with `cellranger mkref` from the filtered GTF and the top-level genome FASTA to support downstream barcode and UMI quantification.

##### S5.5 Primary single-cell preprocessing

We demultiplexed raw sequencing BCL files to FASTQ format with `cellranger mkfastq`, allowing one mismatch during index demultiplexing. We processed the resulting FASTQ files with `cellranger count` against the custom *P. marinus* reference. Cell Ranger performed read alignment, filtering, 10x barcode counting, and UMI quantification with default parameters. We carried out these computations on a Linux workstation.

##### S5.6 Reprocessing and UTR-aware hybrid reference construction

For the direct comparison, we reprocessed two PBS-injected control whole-blood libraries against both the Ensembl-only reference and the Ensembl+Evo2 hybrid reference. This reprocessing pass was distinct from the original Cell Ranger counting reference above. When archived samples were available only as Cell Ranger BAM outputs, we regenerated sample-specific FASTQs with the 10x `bamtofastq` utility bundled with Cell Ranger v10.0.0. We built the UTR-aware hybrid reference by anchoring to Ensembl gene models and appending only non-overlapping Evo2 models with $S \geq 0.99$. In the final reference, this added 17,244 Evo2 loci to the 13,114-gene Ensembl backbone and yielded a 30,358-gene reference. This design preserves the transcript-boundary information relevant to 10x Chromium 5′ capture, especially transcript starts, first exons, and 5′ UTRs, and avoids read siphoning from CDS-only predictions.

#### S6. Pathway Enrichment (cameraPR)

##### S6.1 Gene universe and ranking

The ranking universe comprised structurally annotated Evo2 models with a valid `human_score`. The ranking statistic was the ESM-2 structural similarity to human (`human_score`).

##### S6.2 Reactome gene sets via msigdbr

We loaded Reactome gene sets with `msigdbr` (C2:CP:REACTOME) and mapped them to lamprey genes through the consensus human-match fields in `evo2_consensus_annotation.csv`. We then tested those lamprey gene sets competitively while retaining a stable human-readable pathway namespace.

##### S6.3 cameraPR analysis

We applied `cameraPR` (limma) to the ranked statistics and lamprey-mapped Reactome sets. P-values were adjusted using BenjaminiHochberg FDR. Only Reactome results are reported in the main text to avoid overly broad GO terms. In the main-text immune panel, we collapse overlapping Reactome labels by shared mapped lamprey driver genes before plotting them. That keeps one repeated two-gene signal from appearing as several independent conserved pathways. On the conserved side, FcERI signaling, CLEC7A/Dectin-1 signaling, downstream TCR signaling, and cross-presentation-related labels are driven chiefly by repeated hits to *PSMC3* and *SEM1*, and the broader antigen-processing sets add *SEC61B* plus a few isolated protein-processing hits. Lamprey lacks the canonical mammalian IgE and Fc$\varepsilon$ receptor genes, so those pathway names reflect associated proteostasis and antigen-processing genes rather than those canonical mammalian genes themselves. On the divergent side, Dectin-2 family signaling is driven by *MUC1*/*MUC21*/*MUC3A*/*MUC7*, interferon signaling by *IFI44L*/*PTPN6*/*TUBA4B*, and interleukin-4/interleukin-13 signaling by *MCL1*/*MUC1*. We therefore interpret these Reactome names as similarity-mapped modules rather than literal one-to-one mammalian pathways.

#### S7. Transcription Factor Conservation

To evaluate regulatory remodeling without relying on broad ontology terms, we analyzed the full lamprey consensus annotation against the curated Lambert/Weirauch human transcription factor census (v1.01; $n = 1{,}639$ TFs). Human TF symbols were matched to lamprey consensus calls using the `human_hit` field when available, with fallback to the consensus `final_name`. DNA-binding-domain family assignments were taken directly from the Lambert reference.

For each matched TF group, we computed the mean vertebrate similarity score,

$$\bar{S}_{Vert} = \text{mean}(S_{Human}, S_{Zebrafish}, S_{ElephantShark}),$$

and the branch-shift statistic,

$$\Delta S = \bar{S}_{Vert} - S_{Octopus}.$$

We classified TF groups with the same bilaterian and vertebrate thresholds described above, then summarized them at both the group and locus levels. This yielded 365 TF groups spanning 820 annotated lamprey loci. We ranked family-level summaries by median $\Delta S$ and innovation fraction so we could separate broad TF families that are simply numerous from families that are specifically vertebrate-biased.

| TF | DBD family | $\bar{S}_{Vert}$ | $S_{Octopus}$ | $\Delta S$ | Origin |
|---|---|---|---|---|---|
| ONECUT2 | CUT/Homeodomain | 0.9792 | 0.9448 | 0.0344 | Innovation |
| UBP1 | Grainyhead | 0.9946 | 0.9653 | 0.0293 | Innovation |
| SATB2 | CUT/Homeodomain | 0.9928 | 0.9653 | 0.0275 | Innovation |
| FOSL1 | bZIP | 0.9757 | 0.9488 | 0.0269 | Innovation |
| POU5F2 | Homeodomain/POU | 0.9790 | 0.9560 | 0.0230 | Innovation |
| NFATC4 | Rel | 0.9772 | 0.9588 | 0.0184 | Innovation |

**Table 4 Supplementary Table S3.** Representative vertebrate-biased transcription factor groups highlighted in the main text. Values are taken from the ranked transcription-factor conservation analysis and report mean vertebrate similarity ($\bar{S}_{Vert}$), octopus similarity ($S_{Octopus}$), and the branch-shift statistic $\Delta S = \bar{S}_{Vert} - S_{Octopus}$.

#### S8. Hard-Masked Elephant Shark Rescue Benchmark

To benchmark the full hybrid logic on a vertebrate genome with known truth, we used the Ensembl elephant shark annotation (*Callorhinchus milii*, assembly 6.1.3, release 113) as the accepted reference and the repeat-masked elephant shark genome as the degraded input sequence. The benchmark was carried out in two stages.

| Curated module | Resolved genes used for scoring |
| --- | --- |
| T-like identity | *PTPRC*, *LRRC17*, *RGS8-2*, *LYN* |
| B-like identity | *LRRC15-1*, *A0A8Q3WLD3*, *IGSF6-2*, *LRRC17-2* |
| Iron handling | *NCOA4*, *FTH1*, *FTL-1*, *FTL-3*, *FTL-4* |
| Complement / innate | *CFI*, *C1QB*, *C1R*, *TLR4*, *TLR3-2* |
| Stress response | *mif*, *prdx1*, *prdx5*, *hspa5*, *hsp90aa1.2* |

**Table 5 Supplementary Table S4.** Gene sets used to score the curated immune modules in main-text Fig. 5F. These modules were defined on the resolved hybrid-reference gene names present in the reprocessed immune atlas. The matching machine-readable export is written to `RESULTS/tables/supplementary_table_s4_curated_module_gene_membership.csv`.

#### S8.1 Raw caller-level benchmark

We ran BRAKER2 and Evo2 on the hard-masked elephant shark genome and compared their predicted exon intervals to the Ensembl truth annotation using strand-aware interval overlap. At exon resolution, sensitivity is the fraction of Ensembl truth exons overlapped on the correct strand by at least one predicted exon, precision is the fraction of predicted exons that overlap a truth exon on the correct strand, and F1 is the harmonic mean of the two. This provides the raw component-level benchmark reported in the main text.

#### S8.2 Withheld-reference rescue benchmark

To mirror the lamprey strategy rather than comparing raw callers alone, we created a degraded Ensembl backbone by deterministically withholding 10% of elephant shark protein-coding genes. Withholding was performed at the gene level using a stable hash of `gene_id`, yielding 1,932 withheld genes out of 19,320 eligible protein-coding genes and leaving a reduced backbone of 17,388 genes. Because the split is deterministic, the benchmark is exactly reproducible.

Masked Evo2 predictions were post-processed into transcript models, translated with the same robust ORF-selection procedure used in the lamprey pipeline, and embedded with ESM-2. Each Evo2 protein was then matched to the elephant shark truth proteome in structural space using FAISS cosine search. We evaluated a small grid spanning permissive to stringent structural thresholds, including the permissive setting reported in the main text ($S \geq 0.95$) and the strict benchmark setting ($S \geq 0.9745$), together with minimum protein lengths of 30, 50, and 100 aa. For each setting, we retained only Evo2 models that passed the structural threshold, met the minimum protein length criterion, and did not overlap the reduced Ensembl backbone. These surviving models were then treated as candidate rescue additions.

#### S8.3 Rescue metrics

We report three complementary statistics for each threshold combination:

- **Withheld-gene recall**: fraction of withheld Ensembl genes overlapped by at least one non-overlapping Evo2 rescue model on the correct strand.
- **Added-model hit precision**: fraction of added Evo2 models that overlap a withheld Ensembl gene on the correct strand.
- **Full-truth gene coverage**: fraction of all eligible elephant shark truth genes represented by the reduced backbone alone versus the reduced backbone plus Evo2 rescue additions.

The permissive setting ($S \geq 0.95$, protein length $\geq$ 30 aa) maximized recall but admitted many low-precision additions. We emphasize a stricter setting ($S \geq 0.9745$, protein length $\geq$ 100 aa), which yielded 413 non-overlapping Evo2 additions, rescued 12 of 1,932 withheld genes, and increased total truth-gene coverage from 0.9005 to 0.9011. On a severely degraded vertebrate genome, Evo2 contributes a measurable but limited rescue signal when it acts as a filtered augmentation layer on top of a trusted transcript backbone.

#### S8.4 Benchmark summary

The elephant shark stress test is harsher than the lamprey production workflow. We hard-masked the genome, degraded the backbone by withholding truth genes, and evaluated Evo2 additions only when they stayed non-overlapping relative to the reduced Ensembl reference. Under those conditions, the benchmark reads best as a lower-bound rescue assay rather than an estimate of final annotation performance. Raw Evo2 predictions remain too noisy to serve as a direct vertebrate annotation backbone, but a filtered subset can recover a small number of genuinely missing genes when layered conservatively onto a trusted reference.

##### S8.5 Coverage summaries for the strict and best merged settings

Supplementary Fig. S1 separates the two full-truth coverage bar summaries that we removed from the main-text opening validation figure to improve print-scale readability. Panel A shows the strict benchmark setting ($S \geq 0.9745$, protein length $\geq$ 100 aa). Panel B shows the best merged setting under the combined rescue objective used in the elephant shark grid search. In both cases, the comparison is the same: reduced Ensembl backbone alone versus reduced backbone plus non-overlapping Evo2 rescue additions. Supplementary Fig. S2 preserves the full precision–recall tradeoff scatter across the same rescue grid.

#### S9. Software Versions

| Software | Version |
|---|---|
| Evo2 | 1.0.0 (Arc Institute) |
| ESM-2 | 2.0.0 (Meta AI) |
| FAISS | 1.7.4 |
| BRAKER2 | 2.1.6 |
| RepeatModeler | 2.0.4 |
| RepeatMasker | 4.1.5 |
| Cell Ranger | 8.0.1, 10.0.0 |
| R | 4.3.1 |
| Python | 3.10.12 |
| PyTorch | 2.1.0 |

**Table 6** Software versions used in this study.

#### S10. Computational Resources

All analyses were performed on the Dartmouth Discovery HPC cluster:

- **Evo2 inference**: 1–32× NVIDIA L40S GPUs (job array), 8 CPU cores per shard, 64GB RAM.
- **ESM-2 embedding**: 1× NVIDIA A100 GPU, 16 CPU cores, 64GB RAM; runtime ∼4 hours.
- **FAISS search**: GPU or CPU (IndexFlatIP), typical runtime < 10 minutes.
- **Cell Ranger count**: 24 CPU cores, 128GB RAM; runtime ∼20 minutes per sample.

### Supplementary Figures

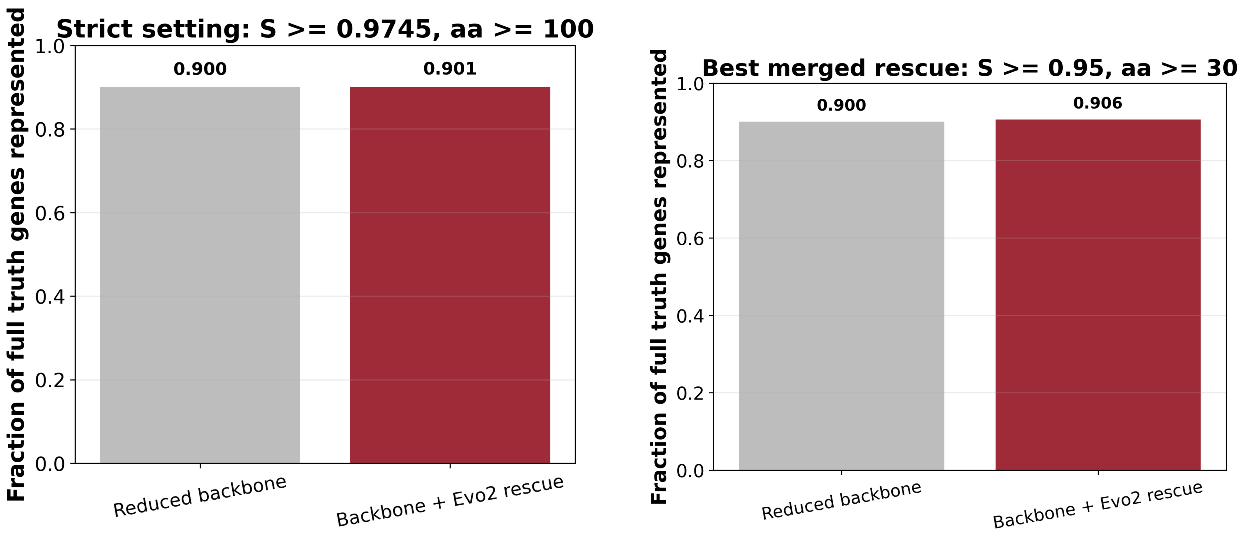


**(A)** Strict benchmark setting

**(B)** Best merged setting

**Fig. 1 Supplementary Fig. S1.** Full-truth coverage summaries for the two elephant shark rescue settings highlighted in the manuscript. (A) Coverage under the strict benchmark setting ($S \geq 0.9745$, protein length $\geq 100$ aa). (B) Coverage under the best merged setting from the elephant shark grid search. In both panels, the reduced Ensembl backbone alone captures most truth genes, and Evo2 rescue additions raise coverage by a small amount.

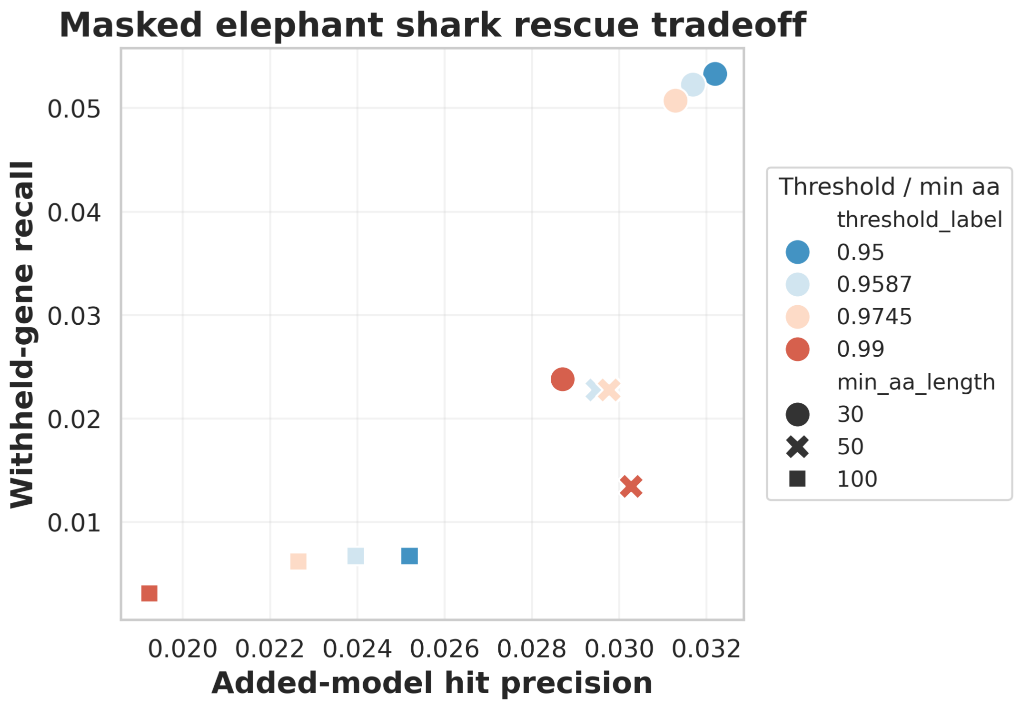


**Fig. 2 Supplementary Fig. S2.** Precision–recall tradeoff across the masked-elephant shark rescue grid. Each point represents one structural-threshold and minimum-length setting used in the rescue benchmark, and point color and marker style encode the threshold and minimum protein-length filter. More permissive settings recover more withheld genes but lower the fraction of added Evo2 models that hit a withheld truth gene. The main-text Fig. 1C keeps only the recall grid so readers see the validation result early without carrying the full tradeoff plot in the opening figure sequence.

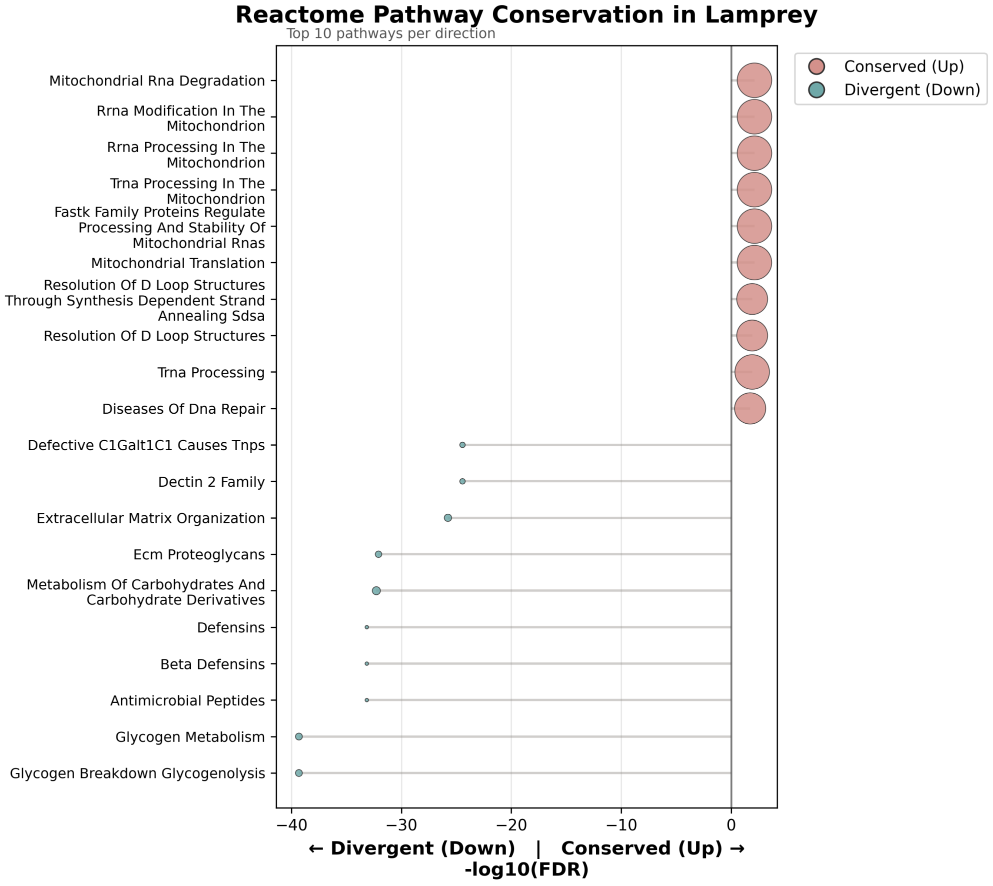


**(A)** Broad Reactome pathway landscape

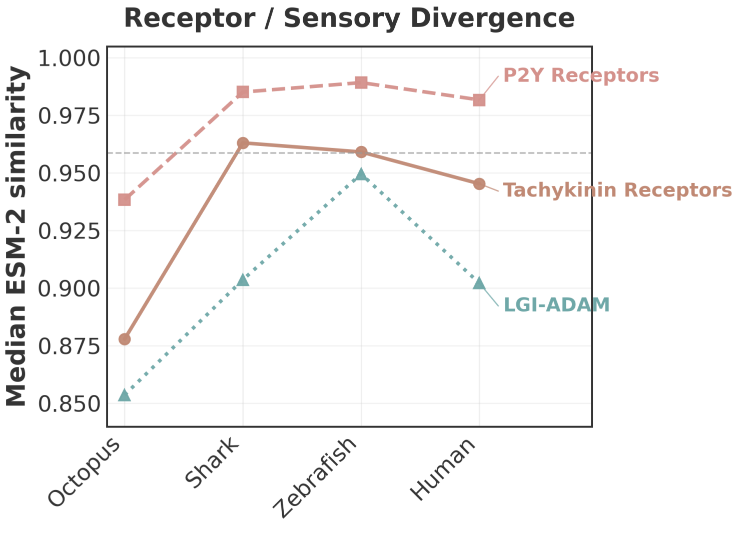


**(B)** Receptor / sensory divergence trajectories

**Fig. 3 Supplementary Fig. S3.** Supplementary pathway-conservation views for the vertebrate-biased structural cohort. (A) Broad Reactome pathway conservation landscape across all Reactome categories. This panel shows the full top-10 up / top-10 down pathway summary, whereas the main-text Fig. 3C collapses overlapping immune Reactome labels into shared lamprey driver modules to emphasize the subset most relevant to the study's biological interpretation. (B) Representative receptor and sensory pathway trajectories, highlighting the strong outgroup drop seen for tachykinin, P2Y, and LGI-ADAM signaling. We moved these trajectories out of main-text Fig. 4 to give the full-width immune directional landscape in Fig. 4C more print-scale space.

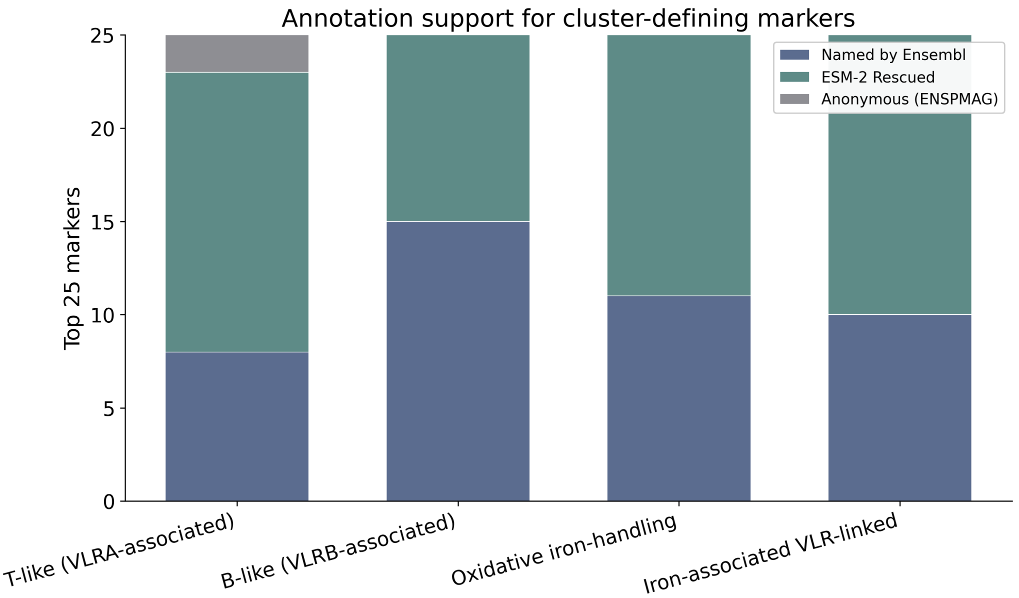


**Fig. 4** **Supplementary Fig. S4.** Annotation support for top marker genes across lamprey immune clusters. The top-marker sets are dominated by named Ensembl genes together with many anonymous structurally supported loci; directly named rescued cross-species assignments are present but do not dominate, and novel Evo2-only models contribute little to these particular cluster-defining markers. This panel therefore serves as a supplementary accounting of annotation provenance rather than as the central single-cell validation panel.

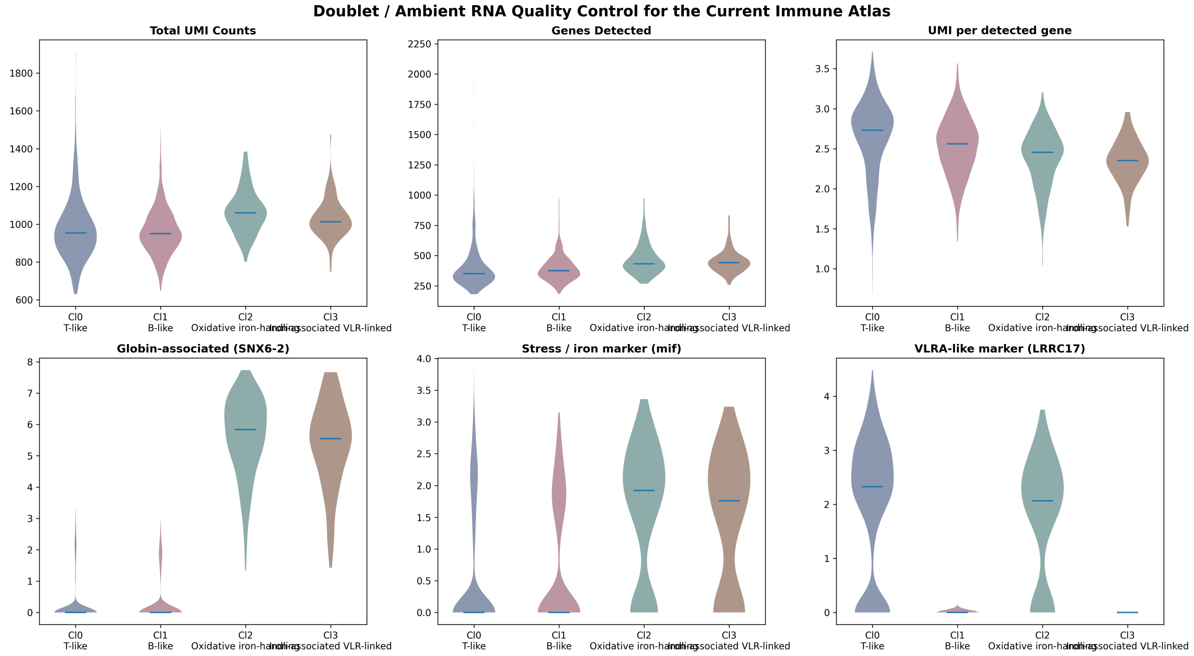


**Fig. 5** **Supplementary Fig. S5.** Doublet and ambient-RNA quality control for the lamprey immune scRNA-seq analysis. The top row shows total UMI counts, genes detected, and UMI-per-detected-gene complexity across the four immune states. The bottom row shows representative globin-linked, stress/iron, and VLRA-associated marker distributions. None of these diagnostics support the interpretation that the oxidative iron-handling and iron-associated VLR-linked populations are dominated by technical doublets or ambient RNA contamination. These diagnostics are consistent with a biological origin of the mixed adaptive-state/iron-recycling signature and argue against a simple clustering artifact.
